# Discovery of prevalent natural *Wolbachia* in *Aedes aegypti* in Metropolitan Manila, Philippines using locally designed primers: bacterial density is influenced by strain and host sex

**DOI:** 10.1101/2022.12.23.521724

**Authors:** Jerica Isabel L. Reyes, Takahiro Suzuki, Yasutsugu Suzuki, Kozo Watanabe

**Affiliations:** Center for Marine Environmental Studies (CMES), Ehime University, Bunkyo-cho 3, Matsuyama, Ehime 790-8577, Japan; Graduate School of Science and Engineering, Ehime University, Bunkyo-cho 3, Matsuyama, Ehime 790-8577, Japan

## Abstract

*Wolbachia* are maternally transmitted bacteria that are utilized for arboviral disease prevention. Cytoplasmic incompatibility (CI) and viral blocking, two characteristics that make *Wolbachia* suitable for vector control, both depend on bacterial infection prevalence and density. Although numerous mosquito species naturally harbor *Wolbachia*, the prevalence of *Wolbachia* in *Aedes aegypti* varies from minimal to absent. The absence of natural *Wolbachia* in *Ae. aegypti* is considered advantageous for novel *Wolbachia* transinfection because interference caused by incompatible *Wolbachia* strains will not occur. In this study, we screened for *Wolbachia* density in 429 individual *Ae. aegypti* collected from Metropolitan Manila, Philippines using locally designed primers to determine whether primer compatibility improves *Wolbachia* detection in naturally infected *Ae. aegypti*. We also investigated the effects of host sex and *Wolbachia* strain on *Wolbachia* density. We found a high *Wolbachia* prevalence for *16S rRNA* (40%) and *wsp* (62%) markers. Relative *Wolbachia* densities ranged from −3.840 to 2.710 for *16S rRNA* and from −4.020 to 1.810 *for wsp*, and these relative densities were higher among male mosquitoes than female mosquitoes. Phylogenetic analysis revealed that most of the *Ae. aegypti* were clustered into supergroup B, with 63% (142/226) of these sequences representing a unique strain referred to here as “*w*AegML,” which exhibited a significantly lower density in *Ae. aegypti* than the other two *Wolbachia* strains detected (*w*AegB/*w*AlbB and *w*AlbA) as well as a higher density in male *Ae. aegypti* than in females. Overall, locally designed primers improved *Wolbachia* detection in naturally infected *Ae. aegypti*. The unique strain *w*AegML occurs at a low density and is related to the use of *w*AlbB in mass-release programs; therefore, it is unlikely to interfere with such programs. Further studies on natural *Wolbachia* infection in *Ae. aegypti* combining methodological and biological factors are warranted to help resolve current conflicts in the field.

## Introduction

The mosquito *Aedes aegypti* is the primary vector for Dengue, Zika, Yellow fever, and Chikungunya viruses, which cause disease in humans [1,2]. To mitigate infections with such viruses, the development of vaccines and the implementation of vector control programs have been prioritized [3]. One vector control method considered a natural and sustainable approach involving the use of the bacterium *Wolbachia*. *Wolbachia* was discovered in *Culex pipiens* [4] and has the ability to induce cytoplasmic incompatibility (CI), wherein gametes fail to produce viable offspring owing to incompatible *Wolbachia* infection [5]. *Wolbachia* also naturally infect and affect arthropods such as *Drosophila spp*. [6–10], *Anopheles spp*. [11,12], and *Aedes albopictus* [13,14]. Different *Wolbachia* strains, i.e., *w*Melpop, *w*Mel, and *w*AlbB, from these natural hosts can exert life-shortening effects [6,10] or confer nutritional benefits [7], efficient maternal transmission [8,15], and antiviral protection [9,16,17]. CI that enables the rapid spread of *Wolbachia* into insect populations coupled with the antiviral effects of *Wolbachia* against arthropod-borne viruses forms the basis for the mass release of *Wolbachia*-transinfected *Ae. aegypti* [18]. Currently, only *D. melanogaster*–derived *w*Mel and *Ae. albopictus*–derived *w*AlbB are used for release programs [18–21] because other *Wolbachia* strains may not exhibit a pathogen-blocking effect [22]. Unlike other host species, conflicting studies on *Ae. aegypti* have mostly revealed the absence of natural *Wolbachia* infection [23–25] but also the occasional presence of *Wolbachia* in *Ae. aegypti* populations without any preceding *Wolbachia* infection. These naturally *Wolbachia*-infected *Ae. aegypti* were collected from the USA [26], Mexico [27], Panama [28], India [29], Malaysia [30], Thailand [31], and Philippines [32,33]. Nevertheless, the majority of the current research focuses on the premise that *Ae. aegypti* do not naturally harbor *Wolbachia*.

Mosquitoes that naturally harbor *Wolbachia* are characterized by consistently high *Wolbachia* infection prevalence, which differentiates them from mosquitoes initially regarded as unnaturally infected [34]. *Culex pipiens, Culex quinquefasciatus, Ae. albopictus, Anopheles moucheti*, and *Anopheles demeilloni* can exhibit 100%*Wolbachia* infection prevalence in contrast to *Anopheles gambiae* and *Ae. aegypti* [11,34,35]. The low and variable natural infection prevalence of *Wolbachia* in *An. Gambiae* and *Ae. aegypti* is 8%–24% and 4.3%–57.4%, respectively, and occurs concurrently with low-density infection that may be due to environmental factors (e.g., temperature and larval contamination) [23,36] or the detection method applied [11,26–28,30]. Nevertheless, both the prevalence and density of *Wolbachia* affect its efficiency as a vector control, given its influence on maternal transmission fidelity and pathogen blocking [11,15,37–43]. Achieving high and stable *Wolbachia* introgression into communities via transinfected *Ae. aegypti* depends on the number of mosquitoes that become infected with the endosymbiont [18,44]. Regarding density, fly hosts such as *Ephestia kuehniella*,carrying closely related natural *Wolbachia* strains (e.g., *w*KueYO and *w*KueTS), and other *Drosophila spp*. exhibit varying CI levels relative to *Wolbachia* density, i.e., high density with high CI [38,40,45–47]. However, one study found that high *w*Mel density in *Drosophila simulans* did not translate into elevated CI but strengthened host immune expression [41]. In *Ae. albopictus*, an increase in *w*AlbB and *w*Melpop-CLA densities reduced Dengue Virus 2 replication *in vitro* [16,48]. Likewise, *Ae. aegypti* mosquito line transinfected with wAlbB inhibited Dengue replication and transmission associated with high immunity and longevity [49].

Bacterial strain and host sex have been linked to *Wolbachia* density. Indeed, different *Wolbachia* strains exhibit not only various densities in hosts [14,50] but also various viral inhibition [9,51] and CI [52] strengths. For instance, *w*AlbB resides in *Ae. albopictus* at a higher density than *w*AlbA [14,50]. In *D. simulans*, only the *Wolbachia* strains *w*Mel, *w*Au, *w*Ri, and *w*No that exist at high density confer viral protection, unlike *w*Ha [9]. In trainsinfected *Ae. aegypti, wMelpop, w*Mel, and *w*AlbB exhibit stable and high-density infection, despite being transferred from their natural hosts, i.e., *D. melanogaster* and *Ae. albopictus* [21,53,54]. Regarding host sex, *Wolbachia* density is reportedly higher in adult females than in males, as observed in *D. simulans* as well as the planthopper species *Laodelphax striatellus* and *Sogatella furcifera* [55,56]. *Wolbachia* in female insect host species were found to be more stable than those in males, which exhibit a decline in *Wolbachia* density with age [38,55,56]. However, in some insects such as *Diaphorina citri*, males exhibit higher *Wolbachia* density than females [39]. In *Ae. albopictus, Wolbachia* density and prevalence vary [57,58]: *w*AlbB density was higher in males than in females, whereas *w*AlbA density was higher in females than in males, and *Wolbachia* density in males continuously decreased with age until *w*AlbA was undetectable [58]; however, a higher prevalence of *w*AlbA and *w*AlbB coinfection was observed in females than in males, in which infection prevalence varied according to the *w*Alb strain [57,58].

Current *Wolbachia* mass release programs focus on *Ae. aegypti*, for which information on prevalence varies and knowledge of the biological factors influencing *Wolbachia* density in naturally infected populations are limited. When *Wolbachia* are naturally present in *Ae. aegypti*, their prevalence is low; thus, it has not been possible to determine their density [28,30,32]. In addition, *Wolbachia* prevalence differs among states and/or cities within a country [59,60]. Previous studies have also used general primers designed based on *Wolbachia* sequences from established natural hosts [27,28,30,32,33]. Therefore, the primers currently used for a particular strain may not accurately detect a wide range of strains [61,62], suggesting that it may be necessary to design primers targeting specific local *Wolbachia* populations. Furthermore, *Wolbachia* density in naturally infected host species is regulated by multiple factors, e.g., host genotype [63], environmental conditions [63], host sex [55,56] and *Wolbachia* strain [14,50]. However, no study has clarified the biological factors that affect *Wolbachia* density in naturally infected *Ae. aegypti*.

The objective of the present study was to determine the biological factors affecting *Wolbachia* density in naturally infected *Ae. aegypti* collected from Metro Manila, Philippines. First, we quantified *Wolbachia* density in 429 individual mosquitoes. Next, we investigated the effects of *Wolbachia* strain and host sex on *Wolbachia* density in natural *Ae. aegypti* populations. We hypothesized that natural *Wolbachia* infection in *Ae. aegypti* is difficult to detect owing to primer incompatibility and that *Wolbachia* density differs depending on *Wolbachia* strain and host sex. The present study provides not only baseline information on the biology underlying natural *Wolbachia* infection in *Ae. aegypti* but also insights into the potential interference of existing *Wolbachia* strains and their density on current mass-release programs [23].

## Methods

### Mosquito sample selection

We used DNA samples extracted from *Ae. aegypti* adult mosquitoes collected in the National Capital Region of the Philippines; these samples were previously used for *Wolbachia* detection via conventional PCR by Carvajal *et al*. [32]. Natural *Wolbachia* was detected in 11.90% (80/672) of these samples [32]; thus, it was considered suitable for validating primer incompatibility. Each individual mosquito was previously screened and processed for DNA extraction as indicated by *Carvajal et al*. [32]. The total DNA of individual mosquitoes was extracted using a Blood and Tissue DNEasy Kit (Qiagen, Hilden, Germany) according to the manufacturer’s protocol with slight modifications [64]. All samples were stored as DNA at −80°C for long-term preservation. Of 672 individual mosquitoes used by Carvajal *et al*. [32], we selected 429 samples based on sufficient volume and DNA concentration for downstream assays.

### *Wolbachia* detection via conventional PCR

Data on conventional PCR results were obtained from the study of Carvajal *et al*. (S1 Table) [32] and used as a baseline reference in the present study in relation to *Wolbachia* prevalence and density detected with newly designed primers. Briefly, Carvajal *et al*. [32] used two known markers targeting the *16S rRNA* gene, which has a slow evolutionary rate, and another marker targeting the highly variable *Wolbachia* surface protein (*wsp*), which is suitable for fine-scale phylogenetic analysis [65]. The sequences of *16S rRNA* and *wsp Wolbachia-specific* primers were as follows: *Wspecf* (5’-GAA GAT AAT GAC GGT ACT CAC-3’) and *Wspecr* (5’-AGC TTC GAG TGA AAC CAA TTC-3’) [61]; *wsp 81F* (5’-TGG TCC AAT AAG TGA TGA AGA AAC-3’) and *wsp 691R* (5’-AAA AAT TAA ACG CTA CTC CA-3’) [65]. PCR amplifications were conducted according to the method of Carvajal *et al*. [32]. Positive *Wolbachia* infection was based on two successful amplifications of both molecular markers.

### Primer design for *wsp* based on local *wsp* sequences

Most *wsp* primers designed for *Wolbachia* detection are strain-specific [14,65]. However, given the rarity of natural *Wolbachia* infection in *Ae. aegypti*, we considered primer incompatibility due to either the presence of new unidentified strains and/or the fast-evolving nature of *wsp*. Thus, we developed new primers specific for our local samples as follows: First, we obtained 118 *wsp* sequences from the *Ae. aegypti* samples of Caravajal *et al*. [32] (GenBank popset 1712729902). Next, a multiple sequence alignment was performed using MUSCLE, and the results were visualized in Codon Code Aligner version 1.2.4 (https://www.codoncode.com/aligner/). The consensus sequence produced from the alignment was then inputted into Primer-BLAST [66] to design *wsp* primers targeting the *Ae. aegypti* samples. Primer-BLAST generated five primer pairs (S2 Table), which were first validated via conventional PCR using a known *Cx. quinquefasciatus* positive sample. Among the primer pairs, primers *wsp* 01 and *wsp* 05 were selected for further optimization, given that they exhibited the correct band size of target markers without nonspecific binding in the sample (S1 Fig). To select the most suitable *wsp* primer pair for downstream analysis, we determined the optimized annealing temperature and primer concentration for both pairs (S2 Fig). Finally, we selected *wsp* 05, given that its PCR efficiency (S3 Fig) fell within the standard MIQE guideline of ≥90% [67].

### Natural *Wolbachia* infection validation using TaqMan qPCR and strain identification via sequencing

To quantify *Wolbachia* density, TaqMan qPCR targeting both *16S rRNA* and *wsp* was conducted. *Wolbachia* quantification was performed using a well-established primer set targeting the *16S rRNA* marker (16SF 5’-AGT GAA GA A GGC CTT TGG G-3’; 16SR 5’-CAC GGA GTT AGC CAG GAC TTC-3’) but with a modification in terms of the fluorescent dye of the probe [22]. Instead of LC640, we used TET as the reporter dye and BHQ1 as its quencher (5’TET-CTG TGA GTA CCG TCA TTA TCT TCC TCA CT-BHQ13’). *Wolbachia* was also confirmed using the newly designed *wsp* primers (*wsp*AAML F 5’-AGC ATC TTT TAT GGC TGGT GG-3’; *wsp*AAML R 5’-AAT GCT GCC ACA CTG TTT GC-3’; wsp probe 5’FAM-ACG ACG TTG GTG GTG CAA CAT TTG C-TAMRA3’) with the *Ae. aegypti* ribosomal S17 (*RPS*17) gene as a reference gene (17SF 5’-TCC GTG GTA TCT CCA TCA AGC T-3’; 17SR 5’-CAC TTC CGG CAC GTA GTT GTC-3’; 17S probe 5’HEX-CAG GAG GAG GAA CGT GAG CGC AG-BHQ13’) [68]. In total, 429 individual mosquitoes were screened for the presence of *Wolbachia* using qPCR with a cut-off Cq value of 35, which was based on an initial qPCR experiment consisting of *Cx. quinquefasciatus* samples representing true natural *Wolbachia* infection and three replicates of the no-template control for each target gene. NTCs yielded a Cq value of ≥ 35; thereafter, negative detections were confirmed using gel electrophoresis (S4 Fig). All singleplex PCR reactions were performed in a final volume of 10 μl containing 5 μl of 2x iTaq Universal Probes Supermix (Bio rad) with 0.3 μM *RPS17* primers, 0.2 μM 16S *rRNA* primers, or 0.5 μM *wsp* primers and 0.2 μM, 0.15 μM, or 0.3 μM of their corresponding TaqMan probes, respectively, with nuclease-free water added to reach the final volume. The following thermal profile was used for both *RPS17* and *16S rRNA* with a CFX96 touch deep well real-time PCR detection system (Bio-Rad Tokyo, Japan): (1) polymerase activation at 95°C for 30 s, (2) denaturation at 95°C for 5 s, and (3) annealing/extension at 60°C for 10 s. The *wsp* thermal profile was as follows: (1) polymerase activation at 95.0°C for 2 min, (2) denaturation at 95.0°C for 5 s, and (3) annealing/extension at 58.8°C for 30 s. Each PCR amplification included a *Wolbachia-*infected *Cx. quinquefasciatus* positive control and a no-template control. *Wolbachia* density was expressed as the relative abundance of *wsp* normalized to *RPS17* [69]. Following detection, qPCR-confirmed *wsp*-positive products were cleaned using a mixture of alkaline phosphatase (TaKaRa) and exonuclease I (TaKaRa). The cleaned samples were subjected to Sanger sequencing for strain identification.

### *Wolbachia* phylogeny

*Wolbachia* phylogeny was inferred using the maximum-likelihood criterion. For this analysis, we used *Ae. aegypti* samples in which the *wsp* gene was detected via qPCR (*n* = 267). We also obtained additional *wsp* sequences from other host species, e.g., *Aedes spp., Anopheles spp., Culex spp*. and others indicated as reference sequences (*n* = 511), from NCBI GenBank (Table 1). In total, 768 *wsp* sequences were aligned using MUSCLE in CodonCode Aligner version 1.2.4 (https://www.codoncode.com/aligner/) and then trimmed to a final length of 103 nucleotide bases. Using DNASp version 6.12.03 [70], we obtained 102 haplotypes, and we subjected the representative sequences of each haplotype to phylogenetic analysis. Tree reconstruction was conducted using only *wsp* given the high evolutionary rate of the gene, i.e., its suitability for strain identification [65,71]. IQ-TREE 2 (http://iqtree.org) [72] was used where the appropriate substitution model was first identified through ModelFinder implemented as a function of the software [73], from which TPM2+G4 was selected as the best-fit model. We set the ultrafast bootstrap approximation (UFBoot) in IQ-TREE to 1,000 iterations, the minimum correlation coefficient to 0.99, and the other parameters to their default settings [74]. For visualization and annotation, we used iTOL (https://itol.embl.de/) [75].

**Table 1.** Reference sequences for *Wolbachia* supergroups.

| Supergroup | Group | *Wolbachia* host species | GenBank No. |
| --- | --- | --- | --- |
| A | wMel | *Drosophila melanogaster* | AF020072 |
|  | wAlbA | *Aedes albopictus* | AF020058 |
|  | wMors | *Glossina morsitans* | AF020079 |
|  | wRi | *Drosophila simulans* (Riverside) | AF020070 |
|  | wUni | *Muscidifurax uniraptor* | AF020071 |
|  | wHa | *Drosophila sechellia* | AF020068 |
|  | wPap | *Phlebotomus papatasi* | AF020082 |
| B | wPip | *Culex pipiens* | AF020061 |
|  | wPip | *Culex quinquefasciatus* | AF020060 |
|  | wAlbB | *Aedes albopictus* | AF020059 |
|  | wAegB | *Aedes aegypti* | MF999264 |
|  | wMa | *Drosophila simulans* | AF020069 |
| Outgroup |  | *Brugia pahangi* | AY527207 |

### Statistical analysis

Unpaired t-tests were used to determine statistical differences between the *Wolbachia* densities of male and female mosquitoes as well as those between the densities of samples detected as positive or negative using conventional PCR. To compare the densities between *Wolbachia* strains, one-way ANOVA with Tukey’s multiple comparison test was performed. Statistical calculations were conducted in GraphPad Prism version 9.2.0 for Windows (www.graphpad.com), and *p* values of *<0.05* was considered statistically significant.

## Results

### Validation of natural *Wolbachia* infection in *Ae. aegypti*

Screening of *Wolbachia 16S* rRNA and *wsp* in *Ae. aegypti* revealed an overall prevalence of 40% (172/429) and 62% (267/429) in the mosquito population, respectively (Table 2). *Wolbachia* density was expressed as the relative abundance of the target gene normalized to *RPS17* [69]. Thus, the median relative *Wolbachia* density of *16S rRNA* and *wsp* was −1.990 and −2.090, respectively. The relative *Wolbachia* densities of *16S* rRNA– and *wsp*-positive samples were between −3.840 and 2.170 and −4.020 and 1.810, respectively. Comparing our qPCR results with the conventional PCR results of Carvajal *et al*. [32], we found that 91% (40/44) of the mosquitoes positive for *16S rRNA* in conventional PCR showed the same result in our qPCR, whereas 9% (4/44) of the mosquito samples were detected as negative via qPCR. Of the 385 samples confirmed as negative for *16S rRNA* via conventional PCR, 34% (132/385) was positive according to qPCR, whereas 66% (253/385) was consistent with negative conventional PCR detection. Regarding *wsp*, 100% (54/54) of mosquito samples that were positive according to conventional PCR were also positive in our qPCR. Conventional PCR *wsp*-negative samples (372) exhibited an infection prevalence of 57% positive (213/372) and 43% negative (159/372) detection according to qPCR.

**Table 2.** Infection prevalence of natural *Wolbachia* in *Ae. aegypti* based on conventional PCR (cPCR) and qPCR.

|  | **Wolbachia-positive/total (% positive)** |  |  |  |  |  |
| --- | --- | --- | --- | --- | --- | --- |
|  | **16S rRNA** |  |  | **wsp** |  |  |
|  | **Female** | **Male** | **Total** | **Female** | **Male** | **Total** |
| **cPCR** | 23/429<br>(5.4%) | 21/429<br>(4.9%) | 44/429<br>(10.3%) | 28/429<br>(6.5%) | 29/429<br>(6.8%) | 57/429<br>(13.3%) |
| **qPCR** | 103/429<br>(24.0%) | 69/429<br>(16.0%) | 172/429<br>(40.0%) | 158/429<br>(36.8%) | 109/429<br>(25.4%) | 267/429<br>(62.2%) |

In an attempt to explain the contrasting negative conventional PCR and positive qPCR *Wolbachia* detection results, we compared the relative *Wolbachia* densities between samples found to be either *Wolbachia-*negative or positive via conventional PCR by Carvajal *et al*. [32] (Fig 1). We found a 30-fold higher median relative *Wolbachia* density in *16S rRNA* (median = −0.8250) and *wsp* (median = −0.8550) *Wolbachia-*positive mosquitoes compared with the conventional PCR–negative samples for *16S rRNA* (*p* < 0.001; median = −2.325) and *wsp* (*p* < 0.001; median = −2.250).

**Fig 1.**
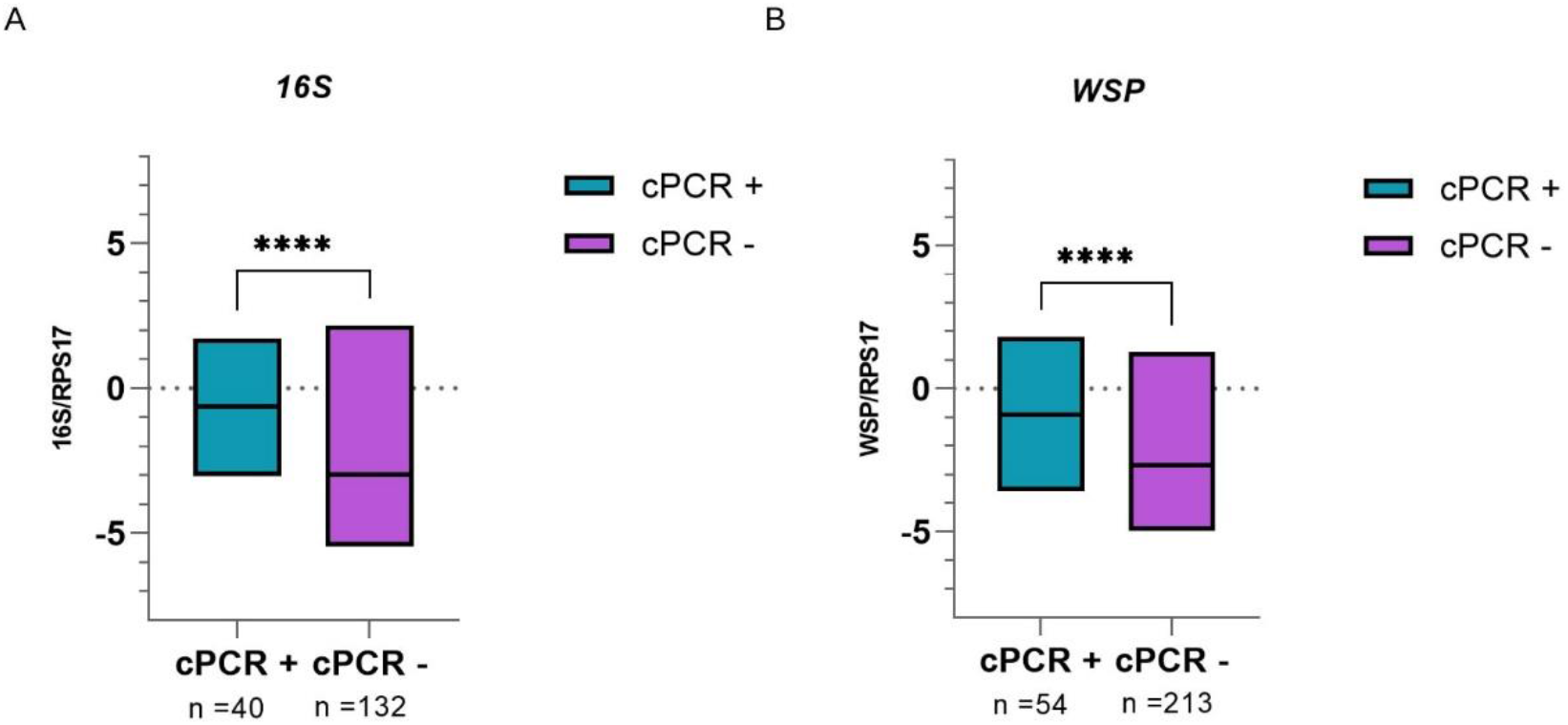
*Wolbachia* density of qPCR-positive *Ae. aegypti* grouped according to conventional PCR results. Individual mosquitoes detected as either positive for *16S rRNA* (n = 172) or *wsp* (n = 267) via qPCR were grouped based on *Wolbachia* detection results according to conventional PCR (cPCR). Relative *Wolbachia* density is expressed as the ratio of the target gene to *RPS17*. Values are shown on a logarithmic scale. (A) Using *16S rRNA* primers, positive and negative samples according to cPCR had a median relative *Wolbachia* density of −0.8250 and −2.325, respectively. (B) Using *wsp* primers, positive and negative samples according to cPCR had a median relative *Wolbachia* density of −0.8550 and −2.250, respectively. **** indicates a significant difference between cPCR-positive and cPCR-negative *Ae. aegypti* at *p < 0.0001*.

### Natural *Wolbachia* density differs according to host sex

We characterized natural *Wolbachia* infection in *Ae. aegypti* according to host sex differences in terms of relative density, finding that *Wolbachia* density differed significantly (*p* < 0.05) between male and female mosquitoes (Fig 2). For both *16S rRNA* and *wsp*, *Ae. aegypti* males exhibited *Wolbachia* densities that were ≥10-fold higher than those of their female counterparts. Regarding the *16S rRNA* marker, male mosquitoes exhibited relative *Wolbachia* densities between −3.570 and 2.170 with a median value of −1.670, whereas females exhibited a relative *Wolbachia* density range from −3.840 to 1.820 and a median value of −2.310. Regarding the *wsp* marker, male mosquitoes exhibited relative *Wolbachia* densities between −3.530 and 1.810 with a median value of −1.880, whereas females exhibited a range from −4.020 to 1.280 and a median value of −2.245. Although *Wolbachia* density was higher in male *Ae. aegypti* than in females, the prevalence of infection was higher among females, regardless of the marker used for detection via qPCR (Table 2).

**Fig 2.**
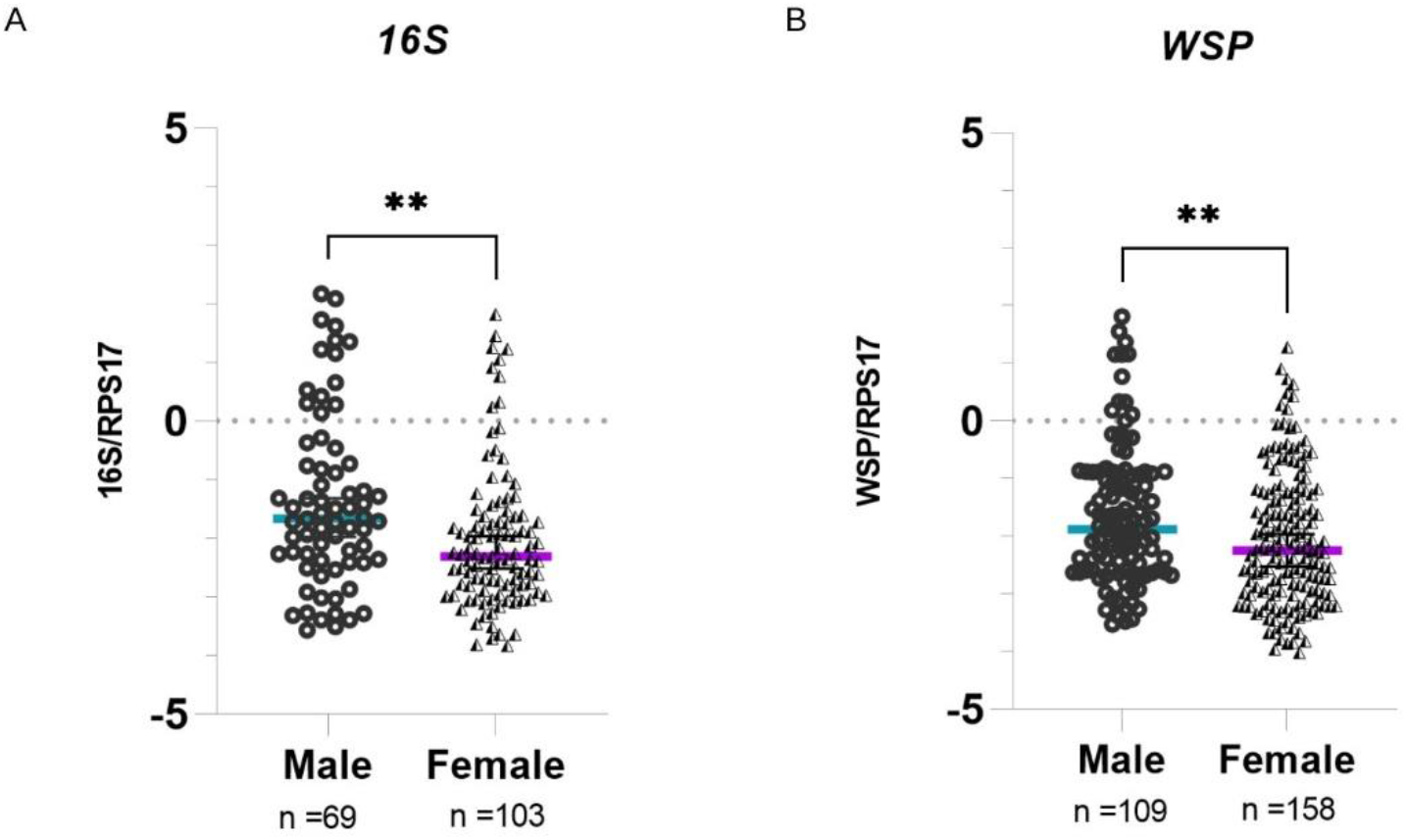
Natural *Wolbachia* infection compared between male and female *Ae. aegypti*. Mosquitoes that were considered *Wolbachia-*positive according to TaqMan qPCR were classified as either male or female. Relative *Wolbachia* density is shown as the ratio of the target gene to *RPS17*. Values are shown on a logarithmic scale. The 95% confidence interval of the median is indicated by the blue and violet lines. Relative densities of (A) *16S rRNA* and (B) *wsp* markers. In both (A) and (B), *Wolbachia* density differed significantly according to host sex. \*\**p < 0.05*. Each data point represents an individual mosquito.

### Phylogenetic analysis of *Wolbachia* strains in *Ae. aegypti*

For phylogenetic analysis, we used 226 *wsp* sequences obtained from 267 *wsp-*positive *Ae. aegypti* samples collected from Metro Manila (AAML). We excluded 41 samples due to low sequencing quality and an inability to repeat sequencing owing to an inadequate volume of these DNA samples. According to the maximum-likelihood phylogenetic tree, 86% of *wsp* sequences in the AAML samples (194/226) were clustered into supergroup B, whereas only 14% of *wsp* sequences (32/226) were clustered into supergroup A (Fig 3). The bootstrap values (≥75%; indicated on the tree branches) supported the divergence of the three clusters among supergroups A and B. Under supergroup A, a clear cluster similar to the strain *w*AlbA was observed. Two *wsp* sequences of *Ae. aegypti* did not exhibit a clear delineation with any *Wolbachia* reference sequence. Numerous weak bootstrap support values (≤74%) were observed in deep nodes within supergroup A, indicating high genetic diversity. Under supergroup B, the *wsp* sequences of *Ae. aegypti* samples were nested within *Wolbachia* cluster *w*AegB/*w*AlbB together with other *wsp* sequences derived from *Ae. albopictus or Culex spp*. However, another cluster was solely composed of *wsp* sequences found in the AAML samples (*n* = 142). For clarity, we hereafter refer to this AAML group (shaded in red in Fig 3) as the “*w*AegML” strain. The two *Wolbachia* strains under supergroup B, i.e., *w*AegB/*w*AlbB and *w*AegML, were rooted from *w*Pip, which was consistent with the grouping previously established by Zhou *et al*. [65]. We also noted that five *wsp* sequences from the *Ae. aegypti* samples formed a distinct cluster that did not fall under any of the supergroups considered; thus, these samples possibly belong to supergroups other than A and B. According to one-way ANOVA (Fig 4), *Wolbachia* density in *Ae. aegypti* carrying the *w*AegML strain was significantly lower than that in *Ae. aegypti* carrying either *w*AegB/*w*AlbB or *w*AlbA. However, *Wolbachia* density did not differ significantly between *Ae. aegypti* carrying *w*AegB/*w*AlbB and *w*AlbA.

**Fig 3.**
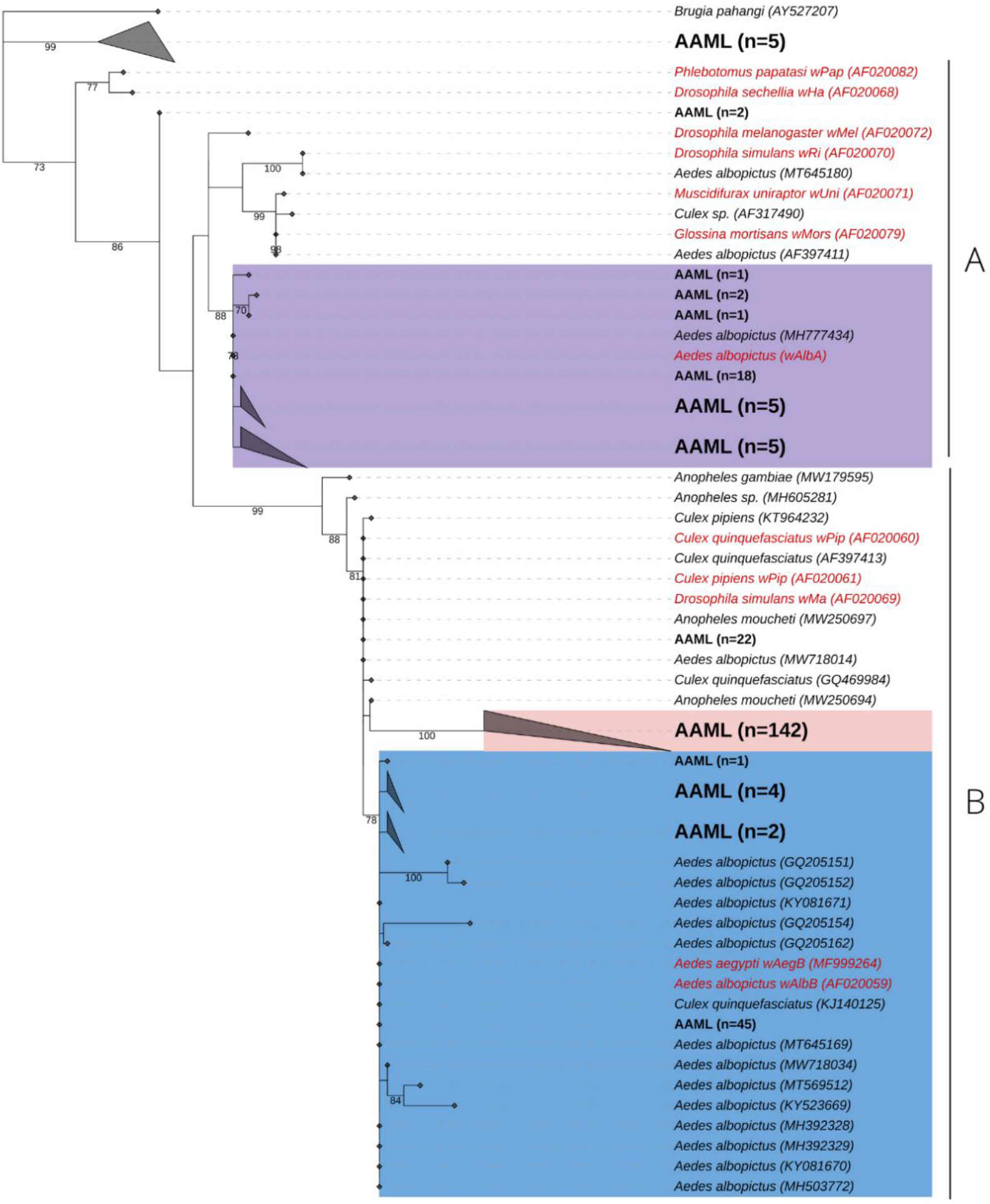
Phylogenetic analysis of *Wolbachia* strains according to *wsp*. Maximum-likelihood tree showing *wsp* sequences from *Ae. aegypti* samples collected in Metro Manila, Philippines (AAML in black bold text) and other *wsp* sequences obtained from other mosquito hosts (data from NCBI Blast; shown in black italicized text). Reference sequences are shown in red with an indication of the corresponding *Wolbachia* strains. Numbers on branches reflect the bootstrap support estimated with 1000 replications. Tree shows three *Wolbachia* strains, namely *w*AlbA (violet), AAML under supergroup B (red), and *w*AlbB/*w*AegB (blue). Scale bar indicates a phylogenetic distance of 0.1 nucleotide substitutions per site.

**Fig 4.**
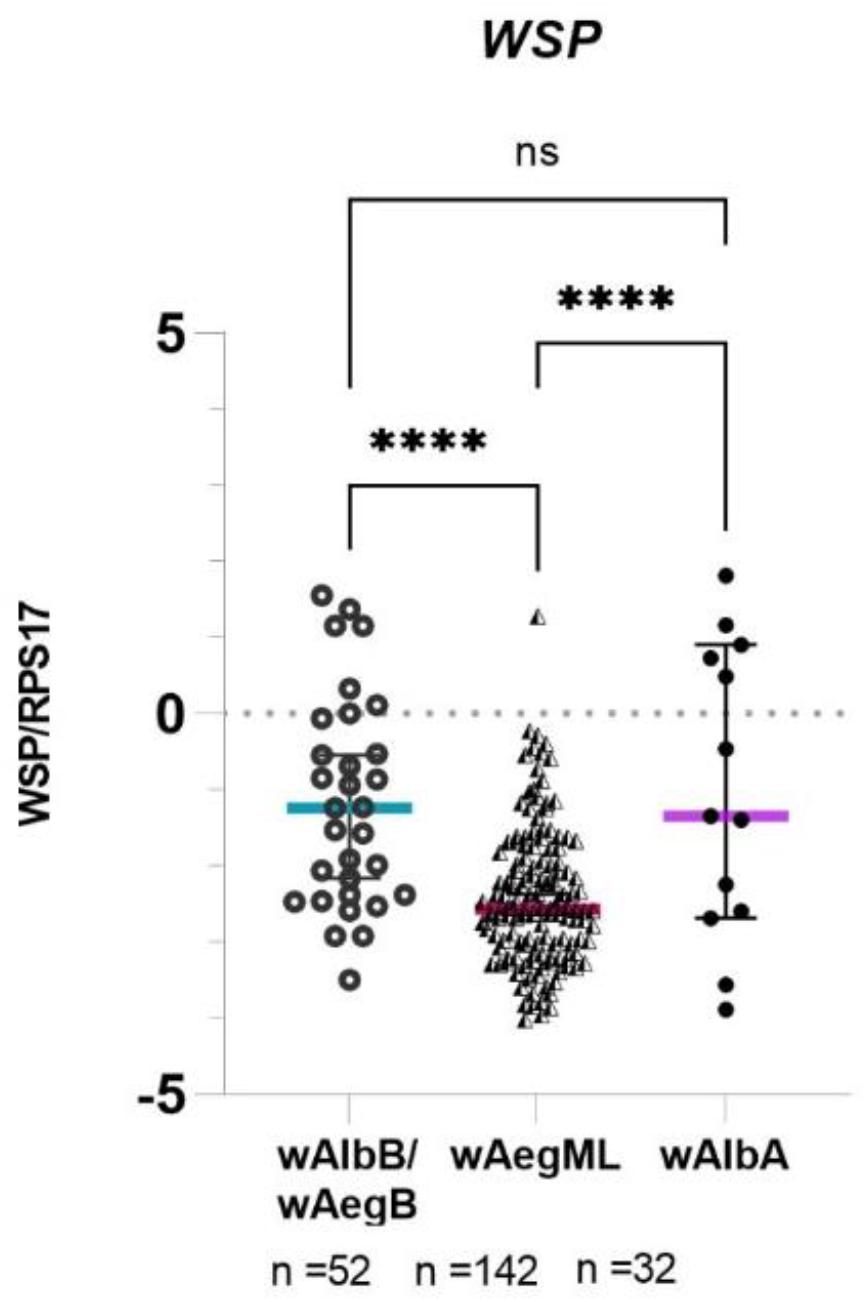
*Wolbachia* density of *Ae. aegypti* according to three identified *Wolbachia* strains. Relative *Wolbachia* density of *Ae. aegypti* in each cluster representing the *Wolbachia* strains *w*AlbB/wAegB, *w*AegML, and *w*AlbA. Relative density was considered the ratio of the target gene to *RPS17*. Values are shown on a logarithmic scale. Median *Wolbachia* relative density is indicated by a colored line. *\*\*\*\*p < 0.05*.

## Discussion

Previous studies on the presence of natural *Wolbachia* in *Ae. aegypti* have resulted in conflicting results and the widely accepted notion that *Ae. aegypti* is uninfected by *Wolbachia* [23–32]. In the present study, we found that relative *Wolbachia* density varied between bacterial strains and according to host sex in natural *Ae. aegypti* populations. The *Wolbachia* strains found naturally infecting *Ae. aegypti* were closely related to the strains found in *Ae. albopictus*, although we identified one unique strain (*w*AegML), which was present at a low density in *Ae. aegypti*. Importantly, using primers designed for local *Wolbachia* populations, we were able to observe a higher prevalence of *Wolbachia* infection. Considering the high genetic diversity among *Wolbachia* strains, the use of locally designed primers may have resulted in a higher detection rate.

### High prevalence of natural *Wolbachia* infection in *Ae. aegypti*

*Wolbachia* is ubiquitous in numerous host species, including lice [76], spider mites [77], planthoppers [55,78], bugs [39], and other insects [38,56,79]. However, this is not the case in mosquito species, in which natural *Wolbachia* infection prevalence is usually high among *Culex spp*. and *Ae. albopictus* but varies according to geographical location among *Anopheles gambiae* (8%–24%) and *Ae. aegypti* (4.3%–58.0%) [34]. In the present study, *Ae. aegypti* collected from Metro Manila, Philippines exhibited an infection prevalence rate of 40.0% and 62.2% when targeting *16S rRNA* and *wsp*, respectively. This finding provides further evidence supporting the presence of natural infection in *Ae. aegypti*, although the prevalence rate in our study was higher than that found in previous studies [26–32]. Given the high prevalence rate of natural *Wolbachia* infection detected here, we speculate that our detection method influenced our results. In a previous study, our laboratory performed conventional PCR–based detection of *Wolbachia* infection in *Ae. aegypti* collected from the current study location, finding a relatively low prevalence considering both *16S rRNA* and *wsp* markers [32]. Likewise, a study conducted by Gloria-Soria *et al*. revealed the absence of *Wolbachia* in 117 *Ae. aegypti* mosquitoes collected from Cebu province, Philippines [25]. In both studies, conventional PCR was used; although the primers used can amplify multiple *Wolbachia* strains, they were not designed using natural *Wolbachia* sequences from *Ae. aegypti* mosquitoes in the local regions. Different *Wolbachia* primers used for PCR assays vary in terms of efficiency and coverage. The performance of 13 *Wolbachia* primer pairs was previously assessed using samples from a wide range of hosts representing supergroups A–F; the results varied, even among primers targeting the same gene, and only two primer sets yielded identical results, with other primers resulting in incorrectly sized amplifications [61]. In the present study, to address the issue of primer incompatibility, we used qPCR and designed primers for *wsp* based on published sequences [32]. Overall, our method led to an improved natural *Wolbachia* detection rate in *Ae. aegypti* owing to primer compatibility.

Notably, the qPCR method requires the use of primers that yield short amplicons (150 bp), which may have contributed to an increase in sensitivity. Additionally, qPCR has a lower limit of detection and higher sensitivity relative to conventional PCR, which could also explain the higher detection rate in the current study [80,81]. When we compared *Wolbachia* density between samples found to be *Wolbachia-*negative or positive using conventional PCR, we found a 30-fold higher median *Wolbachia* density in the *Wolbachia-positive* samples, suggesting that our newly designed *wsp* primers are more compatible with our samples. Notably, we found low *Wolbachia* density that varied according to the bacterial strain (discussed below); therefore, it is likely that both the newly designed primers and qPCR helped increase the natural *Wolbachia* detection rate relative to the detection performance of conventional PCR, resulting in the detection of the higher prevalence rate.

### *Wolbachia* strains found in natural *Ae. aegypti* populations and their densities

The present study adds to the existing evidence that *Wolbachia (wsp)* sequences found in *Ae. aegypti* belong to either supergroup A or B. Most *wsp* sequences found in *Ae. aegypti* collected from Metro Manila, Philippines were categorized under supergroup B. Both supergroups are widespread among arthropods and belong to a single monophyletic lineage [82,83]. Supergroup B is likely to be the dominant supergroup found in naturally infected *Ae. aegypti*, as reported in most previous studies [26,27,29,32]. *Wolbachia* strains under supergroup B usually resides in their hosts at high density [11,50] and exhibit resilience to cyclical heat stress, allowing them to persist in host populations [84].

Interestingly, we detected three clusters representing the *Wolbachia* strains *w*AlbA, *w*AegB/*w*AlbB, and *w*AegML, with the latter strain being unique to our samples. The bacterial density of *w*AegML was significantly lower than that of *w*AlbA or *w*AegB/*w*AlbB, and 62.8% of the *wsp* sequences in our *Ae. aegypti* samples harbored the *w*AegML strain. The *Wolbachia* strains *w*Mel and *w*AlbB are currently deployed in mass release programs as they are maintained stably in *Ae. aegypti* populations [18–21,44]. Pre-existing *Wolbachia* strains in natural *Ae. aegypti* populations could either hamper or strengthen the CI or antiviral effects of *w*Mel and *w*AlbB, depending on strain compatibility [23]. For instance, mating between a host carrying a transinfection of *w*Mel and another host carrying a pre-existing natural *w*AlbB infection will result in reduced or no viable offspring due to the incompatible *Wolbachia* strains [23]. Based on our phylogenetic analysis, it is likely that wAegB and wAegML, which are known to naturally infect *Ae. aegypti*, will be compatible with *w*AlbB used in mass release programs because they are related strains that share the same phyletic lineage, i.e., *w*Pip [29]. In *Cx. pipiens*, different but related *Wolbachia* strains also coexist and are considered compatible [85]. Although the possible effects of these strains on *Wolbachia* release programs are unknown, we can speculate that pre-existing *Wolbachia* infections in natural *Ae. aegypti* populations will have negligible effects that will not interfere with transinfected *Wolbachia* strains used in mass release programs. We also found that the density of *w*AegML was much lower than that of *w*AlbB. Most *Wolbachia-induced* effects are dependent on density [38,40,45–47], and one *Wolbachia* strain tends to become dominant during coinfection with two strains [14,50]. Finally, we found that *w*AegML was the most widespread *Wolbachia* strain in *Ae. aegypti* populations from Metropolitan Manila Philippines.

### Differences in *Wolbachia* density according to *Ae. aegypti* sex

Previous studies have revealed sex-specific *Wolbachia* density differences in natural populations of the planthoppers *S. furcifera* and *L. striatellus* [55] as well as *D. citri* [39], *Drosophila spp*. [56,86], *Cx. pipiens* [87,88], and *Ae. albopictus* [58,89]. We found that natural *Wolbachia* density in *Ae. aegypti* males was higher than that in females, which is consistent with observations in *D. citri* [39] and *Ae. albopictus* [58]. However, such sex-specific variation was only previously observed in *Ae. albopictus* mosquitoes carrying the *w*AlbB strain [58]. Although the biology underlying sex-specific differences in *Wolbachia* density is unknown, biological differences between the host sexes could explain our finding. For instance, female mosquitoes tend to have an expanded microbial composition relative to that of males, resulting in more bacterial competition [90]. Female mosquitoes are hematophagous, and the composition of bacterial microbiota in mosquitoes largely depends on nutrient intake [91,92]. Thus, the digestion process of female mosquitoes may act as a barrier to the survival of some symbionts, including *Wolbachia*. However, others studies have found that females exhibit higher *Wolbachia* densities than males in *S. furcifera, L. striatellus*, [55], *Drosophila spp*. [56,86], *Cx. pipiens* [87,88], and *Ae. albopictus* [58,89]; thus, further investigation on gender-specific effects considering other coexisting factors is warranted. Our study only explored differences of relative *Wolbachia* density between male and female adult mosquitoes. It is important to further characterize how *Wolbachia* density is affected by host sex in different stages of the mosquito life cycle. Determining an absolute *Wolbachia* density rather than a relative one will provide more accurate information. Additionally, supplementing this with microscopic evidence will demonstrate the changes in *Wolbachia* density occurring between male and female mosquitoes.

## Conclusion

Most previous studies on natural *Wolbachia* infection in *Ae. aegypti* have focused on its detection, with the use of multiple methods raised in relation to validating the claim that *Ae. aegypti* is naturally infected [23,25–29,32]. In the present study, we focused on validating the positive detection of *Wolbachia* in *Ae. aegypti* samples previously used by our group [32], which allow us to study the biological *(Wolbachia* strain and host sex) and methodological (primer incompatibility) factors affecting *Wolbachia* prevalence rate and density in natural *Ae. aegypti* populations. Simultaneously studying the methodological and biological factors involved in natural *Wolbachia* infection in *Ae. aegypti* will help resolve current conflicts in the field. Indeed, such studies, including the current study, will help guide existing mass release programs by determining the strains that are less likely to interfere with circulating natural *Wolbachia* strains.

## Acknowledgments

The authors would like to thank Dr. Thaddeus Carvajal of De La Salle University, Department of Biology for his efforts in the field collection of *Ae. aegypti* mosquitoes. We would also like to thank the team leading the Ehime University – De La Salle University International Collaborative Research Laboratory for supporting our study.

## Supporting information

**S1 Table. Conventional PCR (cPCR) and qPCR data on *Ae. aegypti* samples.** The table includes raw data on cPCR and qPCR as well as location and gender of individual mosquitoes.

**S2 Table. Initial list of *wsp* primers and probes generated using PrimerBLAST.** The following primer sets and probes were designed based on the consensus sequence from *Wolbachia* surface *protein* sequences from the study of *Carvajal et al*.

**S1 Fig. Gel electrophoresis results from the initial screening of 5 *wsp* primer pairs using conventional PCR.** Five primer sets generated using PrimerBLAST were validated using conventional PCR. An initial primer concentration of 0.5 μM for each primer was used. One individual sample of *Cx. quinquefasciatus* was used for all primer sets.

**S2 Fig. Gel electrophoresis results of *wsp* AAML01 and *wsp* AAML05 primer sets using different primer concentrations and two different annealing temperatures.** Further primer validation was conducted using wsp AAML01 and wsp AAML05 primer sets. We used two different annealing temperatures 56.9°C and 58.8°C and used two different primer concentrations for each primer set.

**S3 Fig. PCR efficiency of *wsp* AAML05 primer set.** Amplification plot and standard curve of *wsp* AAML05 primer set using 58.8°C annealing temperature and 0.5 μM primer concentration.

**S4 Fig. Representative taqman qPCR amplification plots of each target gene and gel electrophoresis showing no amplification bands using actual *Ae. aegypti* samples.** Amplification plots showing 96-well plate taqman qPCR run of *Ae. aegypti* samples. The representative gel results show selected positive samples (Cq values ≤ 34) and NTC (in green color with Cq values ≥ 35). The gel results show that no amplification was observed in any of the NTCs.

## Notes

### Competing Interest Statement

The authors have declared no competing interest.

